# *In vitro* anti-cancer properties of a commercially available polyherbal nutraceutical (Vernolac) capsule on cancer stem cell-like (NTERA-2 cl.D1) cells

**DOI:** 10.1101/2025.07.15.665027

**Authors:** Poorni Chanuka Rathnayake, Duvinika Chalani Senavirathna, Matheen Muhammadh Milhan, Sandani De Vass Gunawaradane, Nirwani Natasha Seneviratne, Dipun Nirmal Perera, Umapriyatharshini Rajagopalan, Kanishka Sithira Senathilake, Sameera Ranganath Samarakoon

## Abstract

Vernolac is a commercially available polyherbal nutraceutical formulation comprising *Vernonia zeylanica* aerial parts, *Nigella sativa* seeds, *Hemidesmus indica* roots, *Smilax glabra* rhizome, and *Leucas zeylanica* aerial parts. Previous studies have demonstrated anti-cancer activities of phytochemicals derived from these individual plant components. However, the anti-cancer properties of the supercritical CO_2_ extract of Vernolac remain unexplored against cancer stem-like cell populations. The current study is focused on the anti-cancer potential of Vernolac extract on NTERA-2 cl.D1 cancer stem-like model, a human embryonal carcinoma-derived pluripotent cell line. Gas Chromatography-Mass Spectrometry (GC-MS) was performed for phytochemical analysis. Several in vitro assays evaluated the anti-cancer properties of the Vernolac extract. Cytotoxicity was assessed using the Sulforhodamine B assay, and apoptosis induction was determined by Acridine Orange/Ethidium Bromide staining and the caspase 3/7 activity assay. Scratch assay was used to evaluate cell migration, and Reverse Transcriptase quantitative Polymerase Chain Reaction (RT-qPCR) was performed to analyze the expression levels of apoptosis-related genes (*TP53, BIRC5*) and the autophagy-related gene *mTOR*. Further, the free-radical scavenging activity of Vernolac extract was assessed using 2,2-diphenyl-1-picrylhydrazyl and 2,2′-Azino-bis(3-ethylbenzothiazoline-6-sulfonic acid) assay. The reactive oxygen species levels in NTERA-2 cl.D1 cells were quantified using the nitroblue tetrazolium assay. GC-MS analysis revealed 20 phytochemical constituents in the supercritical CO_2_ extract. *In vitro* assay results demonstrated that Vernolac extract exhibits significant anti-proliferative activity in NTERA-2 cl.D1 cells, with an IC_50_ of 41.12 µg/mL at 48 h, while exerting minimal effects on non-cancerous MCF-10A cells (IC_50_ > 1000 μg/mL). Fluorescence microscopy and caspase 3/7 assay showed that Vernolac extract leads to early apoptosis in NTERA-2 cl.D1 cells. RT-qPCR revealed the upregulation of tumor suppressor protein P53 while downregulating *BIRC5* and *mTOR*. Additionally, Vernolac extract inhibited the migration rate of NTERA-2 cl.D1 cells and elevated intracellular reactive oxygen species levels. These findings suggest that the supercritical CO_2_ extract of Vernolac exerts potent anticancer properties against NTERA-2 cl.D1 cancer stem-like model, highlighting its therapeutic potential for targeting cancer stem cells.

## Introduction

Cancer is one of the most common causes of mortality worldwide [1]. Despite various cancer therapies being available, conventional treatments often show limited success due to the challenges, including treatment resistance and tumor recurrence, and adverse side effects [1,2]. These limitations are mainly due to a subpopulation of tumor-initiating cells known as cancer stem cells (CSCs), which possess self-renewal and multi-lineage differentiation capabilities, making them resistant to many conventional therapies, thereby sustaining tumor heterogeneity and recurrence [3–6]. Different treatment strategies are being investigated to overcome the limitations of conventional treatment options. Natural pharmaceuticals are a treatment option currently being investigated for their favorable properties, such as fewer side effects, diverse modes of action, and reduced potential of treatment resistance [7–12]. Moreover, polyherbal extracts are emerging as potential cancer treatment options, and numerous phytochemicals in traditional polyherbal formulations have been shown to exert anticancer effects, with several constituents targeting CSCs through various molecular pathways [13–15]. The NTERA-2 cl.D1 cell line, a well-characterized human pluripotent embryonal carcinoma cell line, is frequently used as a model to study CSC-like model phenotype behavior and evaluate therapeutic responses, serving as a validated *in vitro* platform for evaluating novel anticancer agents targeting stemness-associated pathways [14,16].

Commercially available nutraceutical ‘Vernolac’ is coined after the main constituent *Vernonia zeylanica*. Vernolac is formulated using five different herbal plants, including *Vernonia zeylanica* aerial parts, *Nigella sativa* seeds, *Hemidesmus indica* roots, *Smilax glabra* rhizome, and *Leucas zeylanica* aerial parts [8]. Natural phytochemicals in nutraceuticals reveal multiple advantages through their antioxidant, anti-inflammatory, and cancer prevention potentials [17–23]. Alpha-hedrin, Thymoquinone, and Vernolactone are some of the phytochemicals present in Vernolac formulation, which possess anticancer, antioxidant, and anti-inflammatory properties [15,24–30]. An *in vitro* study conducted by Abeysinghe et al. [15] and Mendis et al. [22] demonstrated that Vernolactone in *V. zeylanica* triggers the antiproliferative activity and apoptosis in NTERA-2 cl.D1 cells, as well as the triads of breast cancer cells (MDAMB-231, MCF-7, SKBR-3) in a time and dose-dependent manner. In addition, Samarakoon et al. [23], Galhena et al. [25], and Pathiranage et al. [26] have reported that a polyherbal mixture comprised of *N. sativa seeds, H. indicus* roots and *S. glabra* rhizome contain possible anticancer and anti-inflammatory effects.

Although the anticancer effects of individual phytochemicals present in the constituent plants of the Vernolac have been demonstrated in various studies [13,15,22,23,31–34], the integrative anticancer effect of the complete polyherbal formulation extracted using the supercritical CO_2_ method has not yet been evaluated, particularly on CSC-like cell populations. Therefore, the present study was designed to investigate the anticancer potential of the supercritical CO_2_ extract of Vernolac against NTERA-2 cl.D1 cells, to elucidate its efficacy as a potential CSC-targeted anticancer therapeutic.

## Materials and methods

### Preparation of Vernolac extract

A polyherbal formulation comprised of *V. zeylanica* aerial parts, *N. sativa* seeds, *H. indica* roots, *S. glabra* rhizome, and *L. zeylanica* aerial parts was extracted using the supercritical fluid extraction method. Plant materials were collected in September 2024 (Batch No. V24085). The extraction was performed using supercritical carbon dioxide (CO_2_) at 42°C, at 30 MPa pressure and the extracts were collected at a CO_2_ flow rate of 4 mL/min for 1 h.

### Cell culture

All the cell lines were cultured and maintained according to American Type Culture Collection (ATCC) guidelines. NTERA-2 cl.D1 (ATCC CRL-1973) cells were cultured in Dulbecco’s Modified Eagle medium (DMEM), and MCF-10A cells were cultured in Mammary Epithelial Cell Basal Medium (MEBM). Both the media types were supplemented with 10%, fetal bovine serum (FBS), 50 U/mL penicillin, and 50 µg/mL streptomycin. Cells were maintained in a humidified incubator with 5% CO_2_ at 37°C.

### Determination of cell viability by SRB assay

Briefly, cells were cultured in 96-well cell culture plates at a density of 7.5×10^3^cells/well in 200 µL of medium for 24h. Cells were treated with varying concentrations of Vernolac extract (6.25, 12.5, 25, 50, 100, 200, and 400 µg/mL) with < 0.1 % DMSO and incubated for 24 and 48h. After incubation, sulforhodamine B (SRB) assay was performed as described in Rajagopalan et al. [35] with minor modifications. Absorbance was measured at 540 nm using a Synergy HT microplate reader, BioTek Instruments, USA.

### Fluorescence microscopic analysis

As described by Abeysinghe et al. [12], NTERA-2 cl.D1 cells (2 × 10^5^ cells/mL) were cultured on a cell culture-treated coverslip and incubated for 24h. The cells were treated with different doses of Vernolac (5.0, 10, and 20 µg/mL) and incubated for 24h. Paclitaxel (1.0, 2.0, and 4.0 µg/mL) was used as a positive control. At 24h post-treatment, cells were washed with ice-cold PBS and fixed with 4% formaldehyde solution in the dark for 10 minutes. Cells were stained using Acridine orange and Ethidium bromide (AO\EB) solution (20 µL). Excess stains were removed with PBS (1 mL); cells were air-dried prior to mounting with anti-fading solution. The stained cells were observed under the fluorescence microscope (BX51 TRF, Olympus Corporation, Tokyo, Japan).

### Caspase 3/7 luminescence assay

NTERA-2 cl.D1 cells were cultured in a cell culture-treated 96-well plate (20,000 cells/ well in 100 μL of media) and incubated for 24h. Cells were treated with different concentrations of Vernolac extract (2.5, 5.0, 10, and 20 µg/mL) and incubated for 24h. Caspase 3/7 Glo^®^ reagent (100 μL) was added to each treated well and incubated for 1h in the dark condition at 37°C as described by Ediriweera et al. [36]. After incubation and 15 minutes in the shaker, Luminescence was measured using a microplate reader (Synergy HT, Biotek Instruments, USA).

### Nitrobluetetrazolium Reactive Oxygen Species (NBT-ROS) assay

NTERA-2 cl.D1 cancer cells (5 × 10^4^ cells/well) were cultured in a 96-well plate using DMEM growth medium with 10% FBS and incubated for 24h. The extract was treated with Vernolac extract 2.5, 5.0, 10, and 20 µg/mL and incubated for 24h. At 24h post-incubation, the NBT-ROS assay was performed according to Ediriweera et al. [37]. Using the Synergy HT microplate reader, absorbance was measured at 560 nm, and ROS expression levels were analyzed as percentage relative values of the untreated control.

### 2,2-diphenyl-1-picrylhydrazyl (DPPH) assay

Free-radical scavenging capacity was measured using 2,2-diphenyl-1-picrylhydrazyl (DPPH) reagent according to the method described by Ediriweera et al. [36]. Volume of 50 µL from each concentration of Vernolac extract (0.01, 0.023, 0.046, 0.093, 0.1875, 0.375, 0.75, 1.5, and 3 μg/mL) prepared with DMSO by the double dilution method. was added in 96-well plates. Samples were further diluted with methanol (90 μL/well) and mixed with 60 μL of DPPH solution (2 mg/ mL) in each well in a dark environment. Ascorbic acid was used as a positive control. The plates were incubated at room temperature in the dark for 10 min, and the absorbance was measured at 517 nm using Synergy HT microplate reader. Free-radical scavenging capacity of each concentration was calculated in percentage using the formula (Absorbance _control_ – Absorbance _sample_)/Absorbance _control_ × 100%, where control contains all the reagents used except Vernolac extract.

### 2,2′-Azino-bis(3-ethylbenzothiazoline-6-sulfonic acid) (ABTS) assay

The 2,2′-Azino-bis(3-ethylbenzothiazoline-6-sulfonic acid) (ABTS) radical scavenging activity of Vernolac extract was measured described by num A dilution series of Vernolac extract (7.8125, 15.625, 31.25, 62.5, 125, 250, 500, and 1000 μg/mL) was prepared in PBS prior to the assay. Vernolac (160 μL) and ABTS (40 μL) were added to each well, and the plate was incubated for 10 min at room temperature. Ascorbic acid was used as a positive control. Absorbance of each well was measured at 734nm using a microplate reader. The percentage radical scavenging activity was calculated as (Abcontrol – Absample) /(Abcontrol) × 100.

### Cell migration assay

NTERA-2 cl.D1 cells were seeded in 96-well plates at a density of 2×10^4^/200 µL and incubated up to 100% confluence until a fully confluent cell monolayer was observed. As described by Abeysinghe et al. [12], a vertical scratch was made on the monolayer formed in each well and washed twice with PBS to remove detached cells and debris. Then, the cells were treated with different concentrations of Vernolac extract (5, 10, 20, and 40 µg/mL) and paclitaxel as the positive control in separate wells. Untreated cells were used as controls. Closure of the scratch made was observed by a phase-contrast microscope at every 12h of incubation, and the area of closures was calculated using the recorded images. The rate of migration affected by respective treatments was calculated using the change in the cell-free area (wound) over time.

### Reverse Transcriptase quantitative polymerase chain reaction (RT-qPCR)

NTERA-2 cl.D1 cells (2.5×10^5^ cells/mL) were seeded into 25 cm^2^ cell culture flasks and incubated for 24h in 37 C and 5 % CO2 : 95 % air environment. Then the cells were exposed to different concentrations (10 and 20 µg/mL) of Vernolac extract under the previously mentioned conditions. Total RNA was isolated using PromegaSV Total RNA isolation system, USA (Cat. No: Z3100SV) according to the manufacturer’s instructions. Complementary DNA (cDNA) from isolated total RNA was synthesized using Moloney Murine Leukemia Virus (M-MLV) reverse transcriptase system (Cat.no: M1701, Promega Corporation, Madisons, USA). Gene expression profiling of *TP53, BIRC5*, and *mTOR* was carried out with *GADPH* gene as the internal control using QuantiNova® SYBR® Green PCR Kit (Cat. No. 208052, Qiagen) and the qPCR reaction was carried out with QuantStudio® 5dx Real-Time PCR Systems (Applied Biosystems, Massachusetts, USA) on 96-well PCR plates. *GADPH* was used as the internal control. The relative gene expression analysis was performed using the Livak method, where fold changes were calculated using 2^−ΔΔCt^ equation [38]. Primers used for gene expression profiling are given in Table 1, and the PCR cycle conditions were as follows: initial denaturation at 95°C for 10 min, and amplification in three steps for 40 cycles (denaturation at 95°C for 30 s; annealing at 58°C for 1 min, and extension at 60°C for 30 s). Annealing temperature for gene *mTOR* was maintained at 52°C.

**Table 01:**
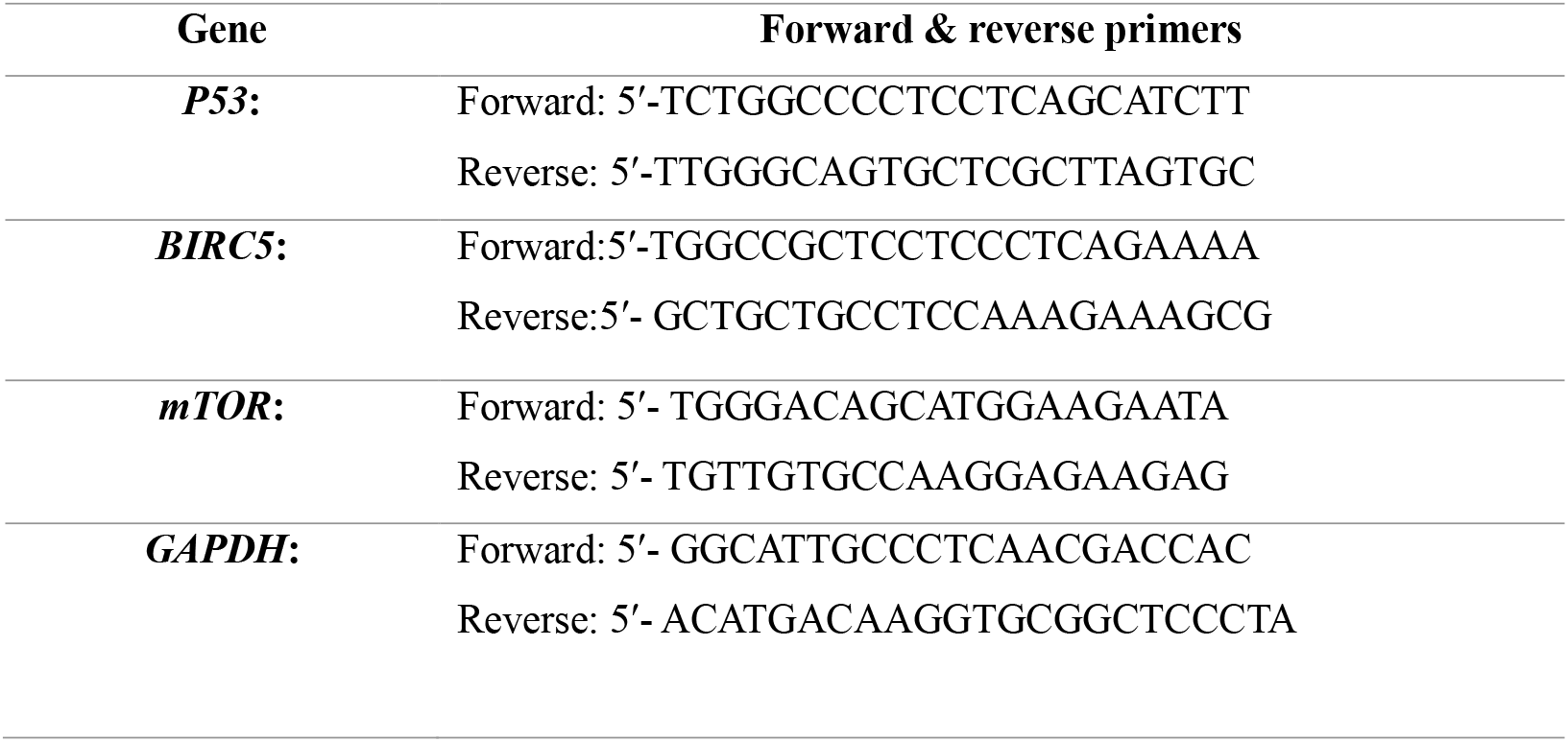
Primers used for expression of *P53, BIRC5, mTOR, GAPDH*.

### Gas chromatography and mass spectroscopic analysis (GC-MS)

GC-MS analysis was conducted on an Agilent 7890 GC system coupled with a 5975 MS detector (Agilent Technologies, Inc., Santa Clara, CA, USA). Separation was achieved on a J&W DB-5 MS capillary column (30.0 m × 250 µm, film thickness 0.25 µm; 5% phenyl methyl siloxane). Ultra-high pure helium (99.999%) served as the carrier gas at a constant flow rate of 1 mL/min. Injections were performed in split mode with a 1:10 ratio using a 1 µL sample volume. Electron ionization was performed at 70 eV ionization energy, while injector and detector temperatures were maintained at 260 °C and 320 °C, respectively. The oven temperature program started at 110 °C (held for 5 min), increased to 280 °C at 20 °C/min (held for 1 min), and then further ramped to 320 °C at 20 °C/min (held for 5 min). Data acquisition and instrument control were managed using MS Solution software. Compound identification was achieved by comparing mass spectra with those in the NIST (National Institute of Standards and Technology) library.

### Statistical analysis

Data analysis was performed using GraphPad Prism 8.0.1 (GraphPad Software Inc., San Diego, CA, USA). Each experiment was triplicated (n=3) for biological triplicates. One-way analysis of variance (ANOVA) was used to compare means across multiple groups in the caspase 3/7 activity assay and NBT-ROS assay, followed by Bonferroni’s post hoc test for multiple pairwise comparisons. For gene expression analysis, one-way ANOVA followed by Dunnett’s post hoc test was performed to compare treatment groups against the control group. Statistical significance was set at P < 0.05.

## Results

### Vernolac extract Characterization

The Vernolac extract yielded 3% (Batch No. V24085). The GC–MS analysis revealed 20 distinct compound peaks. Table 02 summarizes the principal phytochemicals identified in the extract.

**Table 02:**
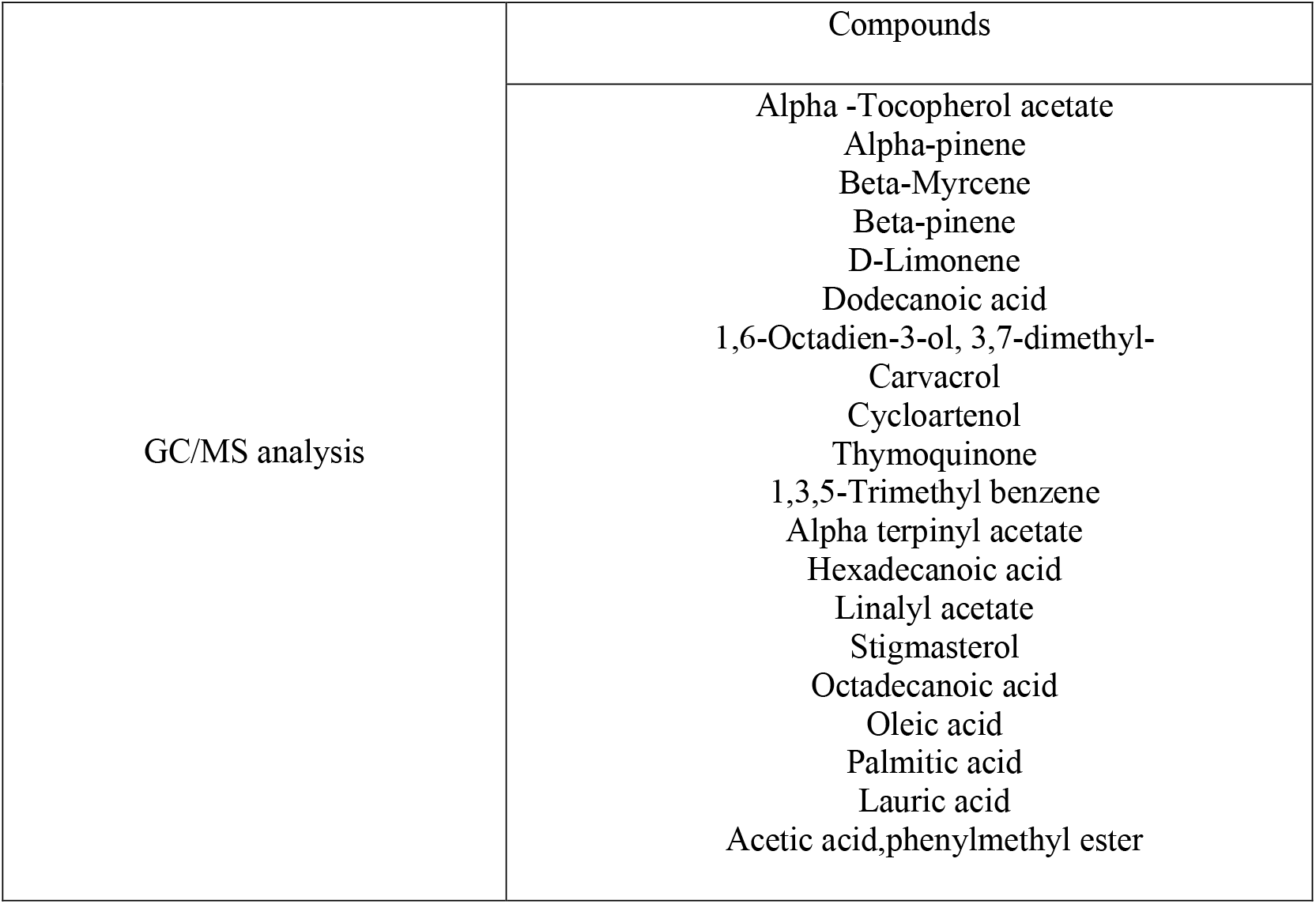
GC/MS analysis of Vernolac Extract.

Notably, several bioactive phytochemicals were identified, including α-Tocopherol acetate, Thymoquinone, Carvacrol, Cycloartenol, Stigmasterol, and Oleic acid, which are known for their antioxidant, anti-inflammatory, and therapeutic properties. Other detected compounds included monoterpenes (α-Pinene, β-Myrcene, β-Pinene, D-Limonene, Linalyl acetate, α-Terpinyl acetate), fatty acids (Lauric acid, Palmitic acid, Hexadecanoic acid, Octadecanoic acid), and additional aromatic or ester derivatives (1,3,5-Trimethyl benzene, Acetic acid phenylmethyl ester).

### Cytotoxicity and cell proliferation assay

As indicated by the results of the SRB assay (Table 03), Vernolac extract exerted a potential to mediate dose- and time-dependent suppression of NTERA-2 cl.D1 cell proliferation. Both Vernolac extract, and Paclitaxel showed a dose and time-dependent anti-proliferative activity against NTERA-2 cl.D1 cells.

**Table 03:** IC_50_ values (μg/mL) of Vernolac extract and Paclitaxel in NTERA-2 cl.D1 and MCF-10 cells.

| Cell type | Vernolac extract (μg/mL) | | Paclitaxel (μg/mL) | |
| --- | --- | --- | --- | --- |
|  | 24h | 48h | 24h | 48h |
| NTERA-2 cl.D1 cells | 64.5 | 41.12 | 4.96 | 1.79 |
| MCF-10A cells | 1075 | 803.5 | 11.90 | 6.70 |

### Fluorescence microscopic observations

Acridine Orange (AO)/Ethidium Bromide (EB) stained NTERA-2 cl.D1 cells were incubated 24h after the treatment with different concentrations of Vernolac extract and Paclitaxel. By employing Acridine orange/Ethidium Bromide staining, clear identification of apoptosis-associated membrane alterations was achieved. The apoptotic changes of the cells were distinguished and visualized by the fluorescence staining, yellow/orange fluorescence indicates early apoptotic cells while red fluorescence indicates late apoptosis, demonstrating disruption of cellular membrane. The induction of apoptosis was increased in Vernolac extract (5, 10 and 20 µg/mL) - and Paclitaxel (1, 2, and 4 µg/mL) treated NTERA-2 cl.D1 cells in a dose-dependent manner (Figure 01).

**Figure 01.**
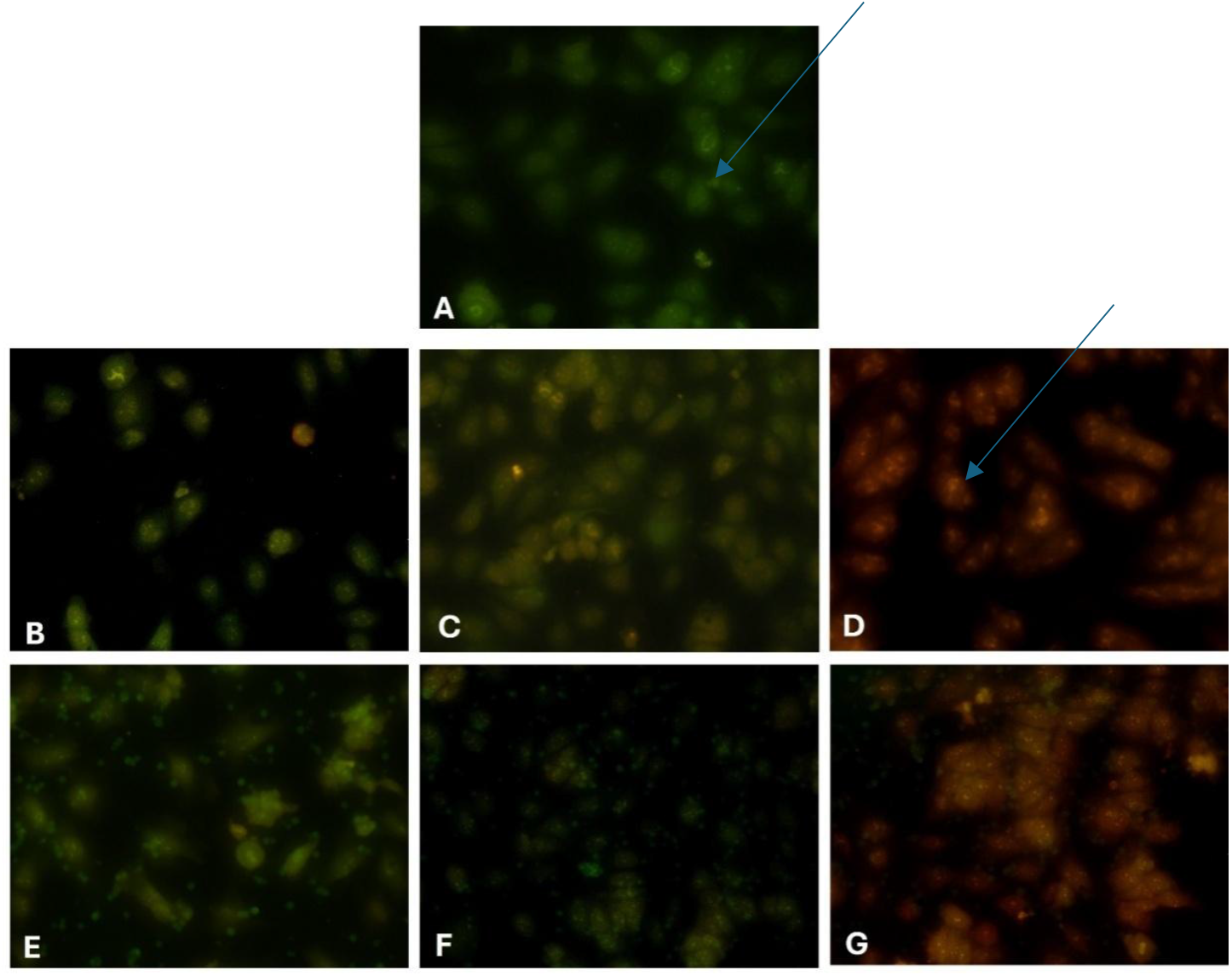
AO/EB stained Fluorescence microscopic observation of NTERA-2 cl.D1 cells under 200X magnification 24h post-treatment incubation. A – Untreated control, B – Paclitaxel (1 µg/mL), C – Paclitaxel (2 µg/mL), D – Paclitaxel (4 µg/mL), E –Vernolac extract (15 µg/mL).F– Vernolac extract - (30 µg/mL), G –Vernolac extract (60 µg/mL).

### Expression of caspase 3 and Caspase 7 activity in NTERA-2 cl.D1 cells

Activities of caspase 3 and 7 were assessed using the Caspase Glo®3/7 assay kit (Promega, USA) according to the manufacturer’s guidelines. Results are presented as the mean ± standard deviation values. Vernolac extract significantly increased the activities of caspase 3 and caspase 7 in NTERA-2 cl.D1 cells at the concentrations of 10 μg/mL (P < 0.001) and 20 μg/mL (P < 0.0001) as shown in Figure 02.

**Figure 02.**
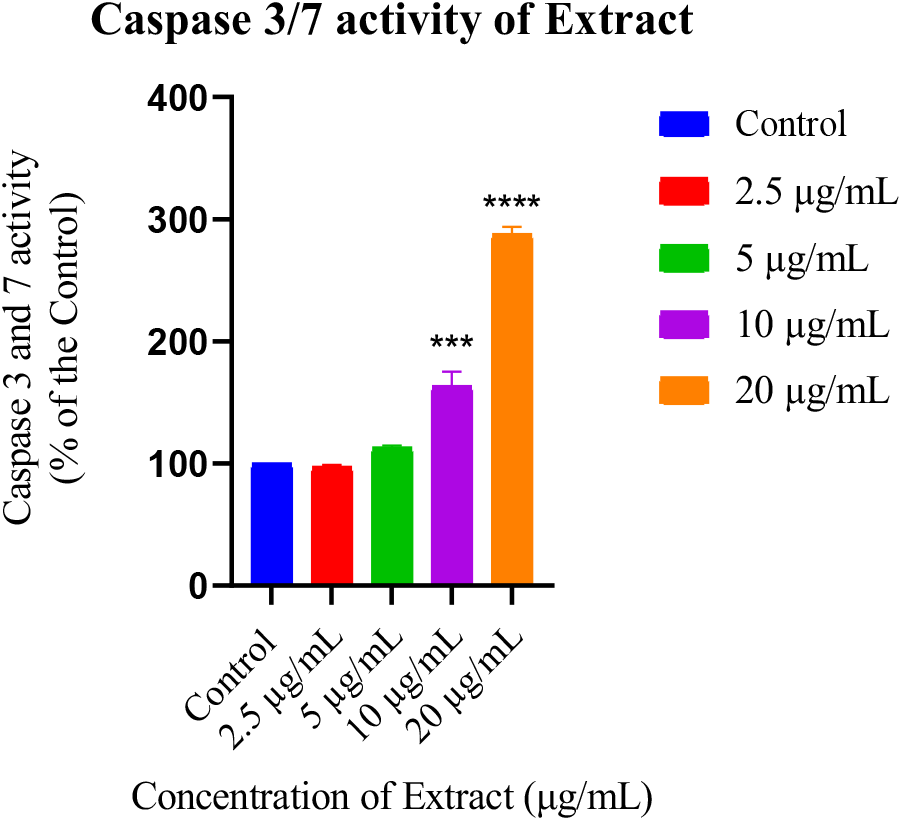
Expression of Caspase 3 and 7 in NTERA-2 cl.D1 cells after 24h post-treatment with Vernolac extract. (Results are indicated as mean ± standard deviation of duplicate determination. *** P < 0.001 and ****P < 0.0001 compared to the control).

### Antioxidant activity of Vernolac Extract

The 2,2-diphenyl-1-picrylhydrazyl (DPPH) assay and 2,2′-Azino-bis(3-ethylbenzothiazoline-6-sulfonic acid) (ABTS) were performed to evaluate the antioxidant activity of Vernolac extract. Ascorbic acid was used as a positive control in both assays. Vernolac extract showed a low free radical scavenging activity (EC_50_ > 1000 μg/mL) in both ABTS and DPPH assays. The obtained EC_50_ values of free radical scavenging activity of Vernolac extract and Ascorbic are summarized in Table 04.

**Table 04:** EC_50_ values of free radical scavenging activity for Vernolac extract and Ascorbic acid.

| Sample | $EC_{50}$ values ($\mu\text{g/mL}$) |
| --- | --- |
| Vernolac extract | > 1000 |
| Ascorbic acid | 53.678 |

### Effect of the Vernolac extract on ROS levels

NBT-ROS assay was performed to analyze the ROS production in NTERA-2 cl.D1 cells after 24h of Vernolac extract. A P < 0.001 was considered to indicate a strong statistically significant result. As seen in the bar graph (Figure 03), treatment with 10 µg/mL or 20 µg/mL of Vernolac extract caused a major increase in ROS activity compared to the non-treated control group (P < 0.001).

**Figure 03.**
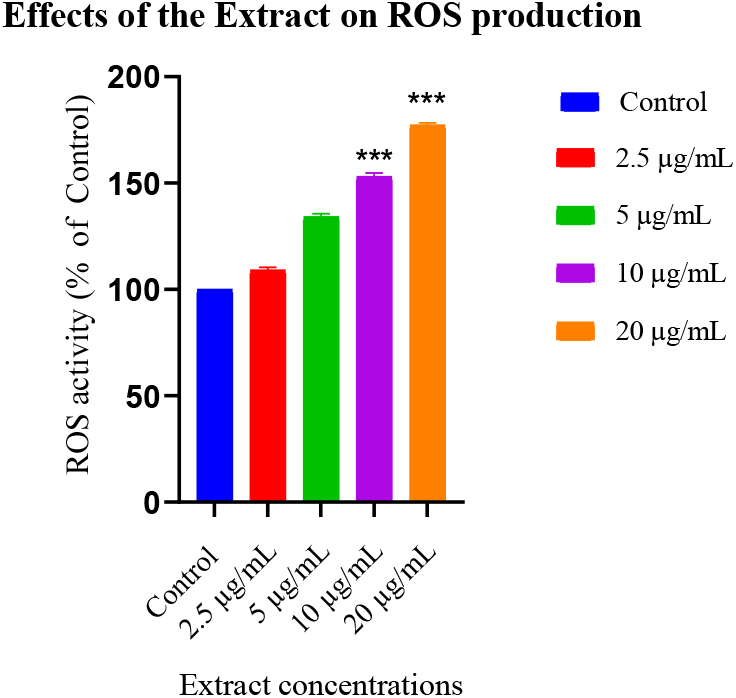
Levels of reactive oxygen species (ROS) detected in NTERA-2 cl.D1 cells after 24h incubation with Vernolac extract treatment under different concentrations (Results are indicated as mean ± Standard deviation. ***P < 0.001 compared to the control).

### Analysis of the wound healing rate of NTERA-2

The NTERA-2 cl.D1 cells were incubated until they created a confluent monolayer at the bottom of the well. An artificial gap was made on the confluent monolayer, and both Vernolac extract and Paclitaxel were treated separately according to a concentration gradient. After 24h of drug treatment, untreated control cells have migrated at a rate of 62.126 μm/h. The cell migration rate of Vernolac extract and Paclitaxel is graphically represented as shown in Figure 04. The morphological representation of cell migration at 0h and after the gap of control closure is shown in Figure 05.

**Figure 04.**
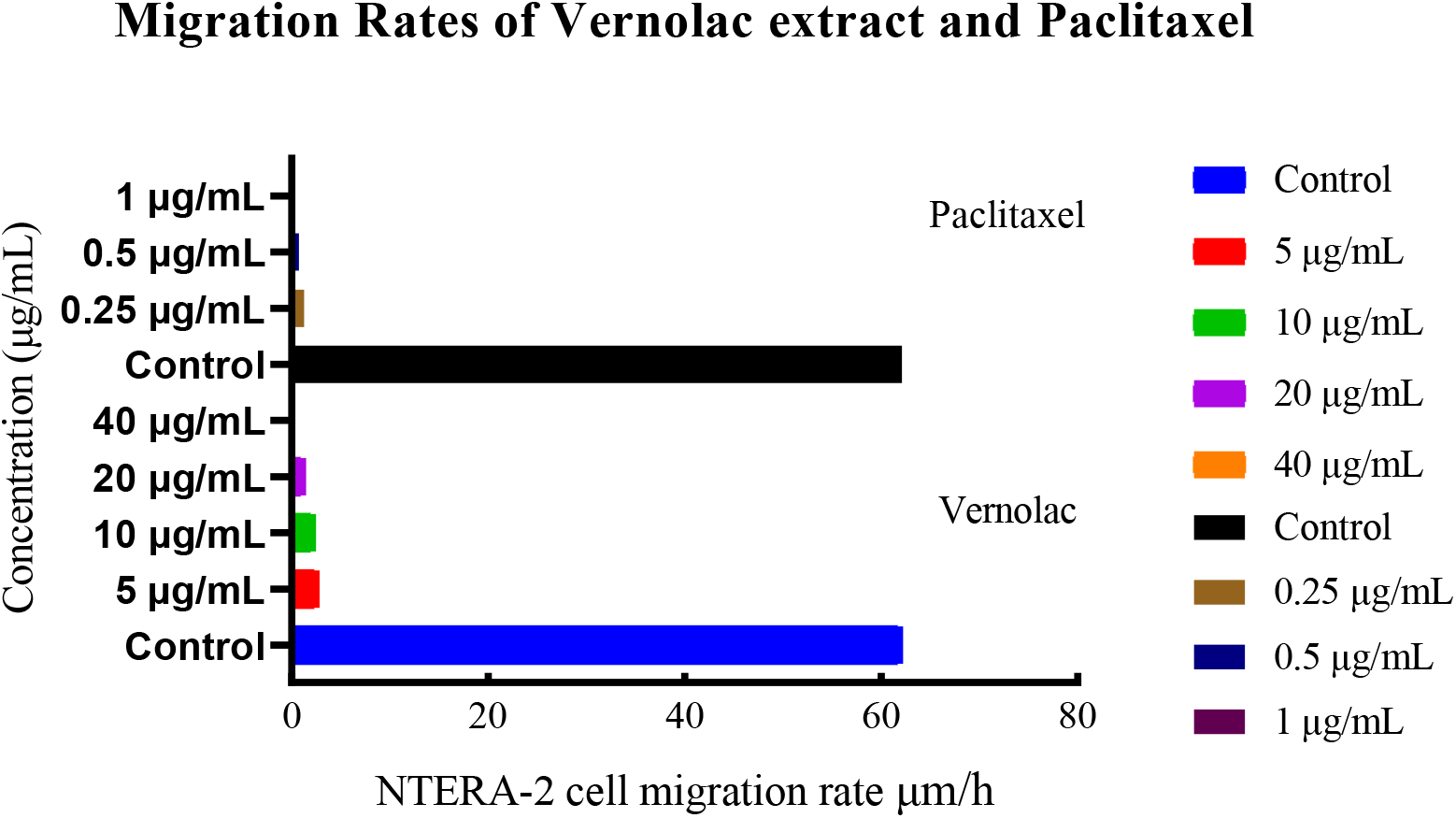
Cell migration rates of Vernolac extract with the concentrations of 5, 10, 20, and 40 µg/mL and Paclitaxel 0.25, 0.5, and 1 µg/mL treated NTERA-2 cl.D1 cells compared to untreated control under different concentrations.

**Figure 05.**
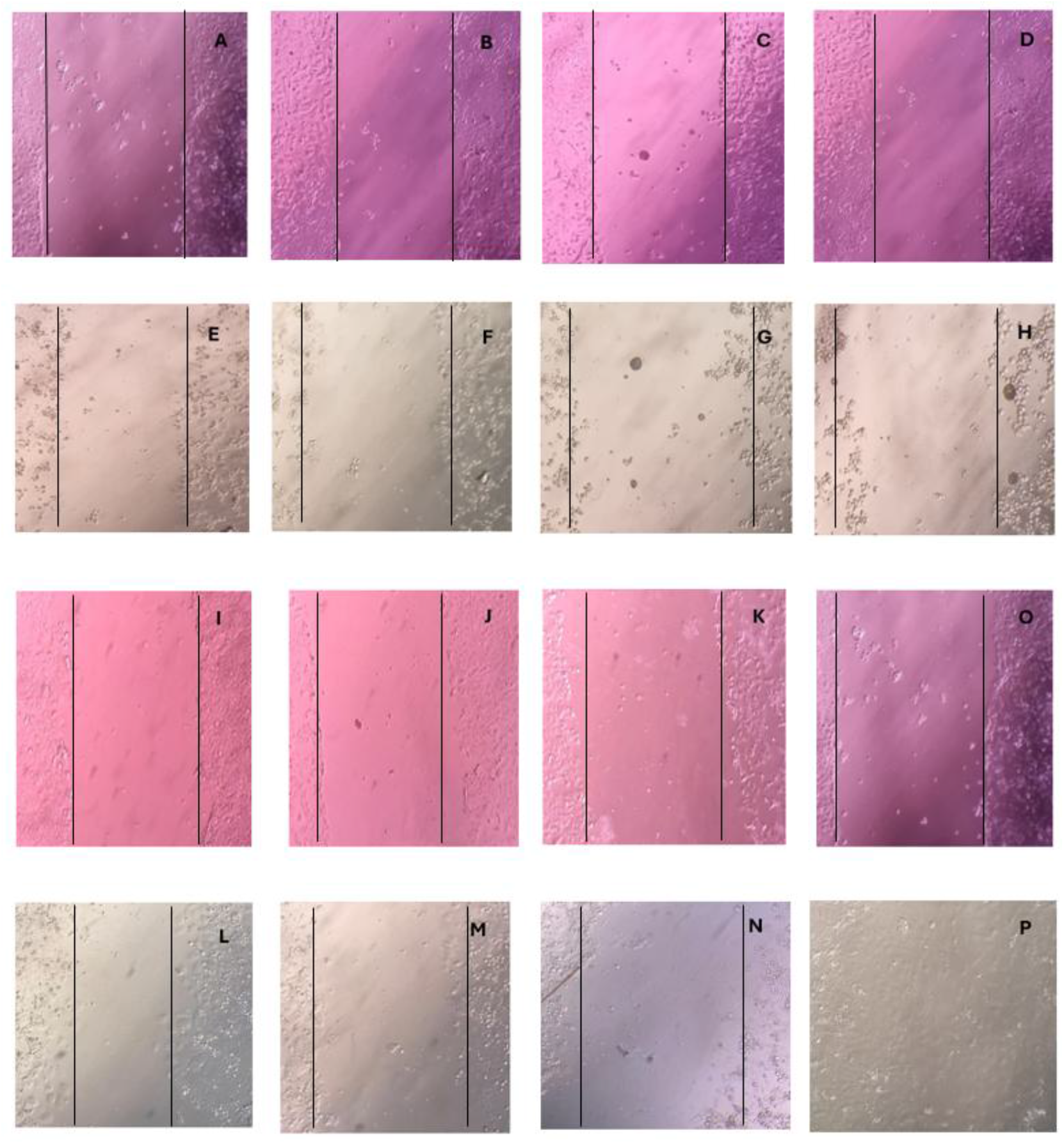
Phase contrast light microscopic images under 10× magnification captured after 0h of creating the gap in NTERA cl.D1 cell monolayer, followed by Vernolac extract with 5, 10, 20, and 40 µg/mL concentrations (A-D), paclitaxel with 0.25, 0.5, and 1 µg/mL concentrations (I–K), and the Untreated control (O). Cell migration after 22h of creating the gap, followed by Vernolac extract (E-H), paclitaxel (L–N), and Untreated control (P).

### Effect of Vernolac extract on the expression of *P53, BIRC5*, and *mTOR*

Results of the RT-qPCR (Figure 06) demonstrated that the Vernolac extract enhanced *TP53* mRNA expression in human NTERA-2 cl.D1 cells in a significant (P < 0.001) dose-dependent manner. Furthermore, treatment with all concentrations of supercritical Vernolac extract and Paclitaxel resulted in a significant downregulation (P < 0.05) of the expression of *mTOR* and *BIRC5*.

**Figure 06.**
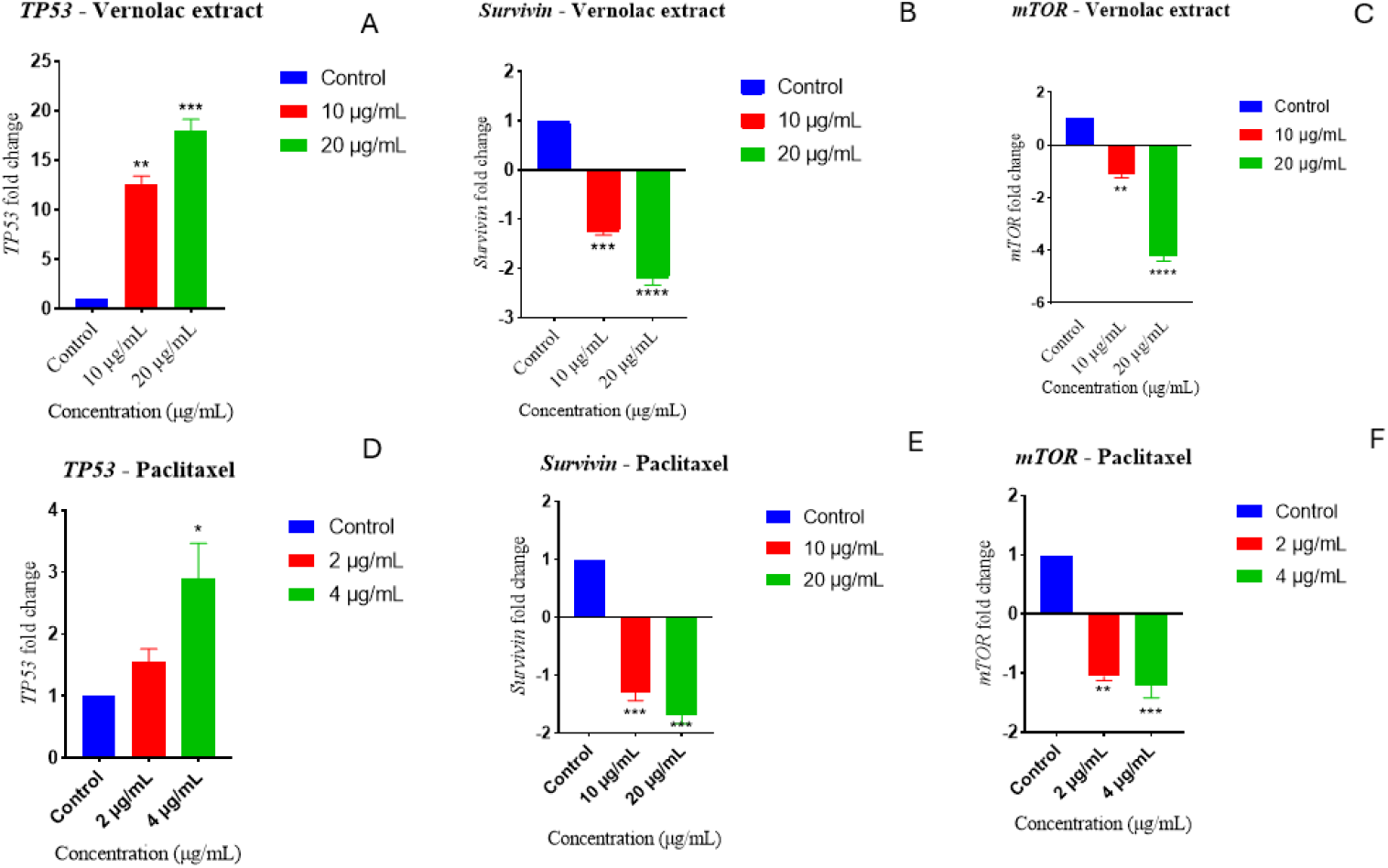
mRNA expression of selected genes on Vernolac extract and Paclitaxel treatment. (A-Vernolac extract on *TP53* expression, B - Vernolac extract on *BIRC5* expression, C - Vernolac extract on *mTOR* expression, D-Paclitaxel on *TP53* expression, E - Vernolac extract on *BIRC5* expression, F - Vernolac extract on *mTOR* expression). (Results are indicated as mean ± standard deviation, duplicate determination. *P < 0.05, **P < 0.01, ***P < 0.001, and ****P < 0.0001. compared to the control).

## Discussion

Previous studies have shown that traditional polyherbal formulation demonstrates significant anti-proliferative activities against several cancer cell lines, including MCF-7, HepG2, and NCI-H292 [23,26,32,39,40]. Sailu et al. [13] and Abeysinghe et al. [15] have elucidated that phytochemicals such as alpha-hederin and Vernolactone present in the constituents of Vernolac have proven anticancer activity against cancer stem-like cells. Chemical analysis using GC/MS identified 20 compounds in all the biological Triplicates of Vernolac extract (Table 02). The present *in vitro* investigation employed supercritical fluid extraction with liquid CO_2_, a superior extraction methodology that preserves thermolabile compounds while eliminating organic solvent residues, to evaluate the anticancer efficacy of Vernolac extract against the pluripotent human testicular embryonal carcinoma (NTERA-2 cl.D1) cell line.

Consistent with these earlier findings, as shown by Abeysinghe et al. [12], the profiles obtained through SRB assay of this study demonstrated dose and time-dependent cytotoxicity against NTERA-2 cl.D1 cells. Vernolac extract showed an anti-proliferative effect against the cells with an IC_50_ value of 41.12 µg/mL after 48h treatment, while demonstrating significantly lower cytotoxicity against non-tumorigenic epithelial MCF-10A cells (IC_50_ value of 1075 µg/mL), suggests selective cytotoxicity towards cancer stem cells (Table 03). ‘Vernolactone’, which is an active compound of the polyherbal isolated from *V. Zeylanica*, has been shown a significant cytotoxicity against NTERA-2 cl.D1 (0.4808 µg/mL at 72h) cells with Paclitaxel [15]. Similarly, Mendis et al. [22], Samarakoon et al. [23] and Pathiranage et al. [26] have also reported that a polyherbal mixture, including several constituents of Vernolac (*N. sativa, H. indicus* and *S. glabra*,) displayed a dose-dependent inhibition of cell proliferation in treated cancer cell lines (MCF-7, MDA-MB-231, HepG2), aligning with the findings.

The fluorescent and phase-contrast microscopic data revealed that Vernolac extract induces dose-dependent apoptosis-related cytomorphological changes in NTERA-2 cl.D1 cells (Figure 01), including cell membrane blebbing and cell shrinkage. Furthermore, compared to the untreated cells (control), Vernolac extract significantly (P < 0.0001) enhanced the enzymatic activities of executioner caspases, caspase 3, and caspase 7 at a 20 µg/mL concentration, indicating activation of the terminal phase of programmed cell death (Figure 01). Concentration gradient for the Caspase 3/7 assay was obtained after several dose optimizations, and depending on the Caspase 3/7 activity results, the concentration of Vernolac extract for the other core assays, including NBT-ROS and RT-qPCR has been decided. These findings align with previous research done by Abeysinghe et al. [15], who reported that Vernolactone significantly (P < 0.0001) upregulated caspase 3/7 activity at both 2 μg/mL and 4 μg/mL concentrations when compared to the untreated NTERA-2 cl.D1 cells (untreated control). A traditional polyherbal mixture formulated using *N. sativa, H. indicus* and *S. glabra* has also demonstrated an elevated caspase 3/7 activation against several cancer cell lines (HepG2, MCF-7, MDA-MB-231, SKBR-3), suggesting a conserved apoptotic mechanism among these phytotherapeutic agents [22,26,32]. Further, extracts of *N. sativa* seed extract demonstrated an elevated caspase 3 activity on the MDA-MB-231 breast cancer cell line.

In this study, the DPPH and ABTS free radical scavenging assays revealed that Vernolac extract contains a low antioxidant activity (EC_50_ value > 1000 μg/mL) compared to the positive control: Ascorbic acid (EC_50_ = 53.678 μg/mL) (Table 03). The obtained result aligns with a previous study demonstrating that Vernolactone possesses low antioxidant activity (free radical scavenging capacity) through the DPPH and ABTS assays [15]. However, multiple investigations have indicated that *N. sativa, H. indicus*, and *S. glabra* extracts have shown free radical scavenging activity through multiple analyses [26,31]. Comprehensive antioxidant profiling using complementary analysis is recommended to fully characterize the antioxidant capacity of Vernolac extract. Despite, extract of Vernolac having a minimal antioxidant activity, a significantly (P < 0.001) elevated ROS level was determined in NTERA-2 cl.D1 at the concentrations of 10 and 20 µg/mL (Figure 03). This pro-oxidant effect in cancer cells, contrasting with limited antioxidant capacity in cell-free systems, suggests a selective modulation of redox homeostasis within the cellular microenvironment, potentially contributing to the observed cytotoxicity [41].

The effect of Vernolac extract on the migration of NTERA-2 cl.D1 cells was evaluated using a wound healing assay. A dose-dependent migration rate (< 3 μm/h across all tested concentrations) (Figure 04), demonstrated minor or no reduction in the wound width throughout the experiment in Vernolac-treated cells compared to the untreated NTERA-2 cl.D1 cells (Figure 05), indicates significantly reduced migratory capacity. Similarly, paclitaxel also restricted cellular migration to rates below 1.5μm/h for all tested concentrations. Low serum concentrations in the cell medium were maintained to ensure the suppression of cell proliferation [42]. This pronounced inhibition of cellular motility indicates impairment of cytoskeletal reorganization and extracellular matrix interaction, critical processes in tumor cell invasiveness and metastatic potential [43].

Moreover, RT-qPCR was performed to evaluate the mRNA expression of several autophagy and apoptosis-related genes (*TP53, BIRC5*, and *mTOR*). The tumor suppressor *TP53* is a key gene that is essential for DNA repair, apoptosis, and triggering of cell cycle arrest, and plays a role in autophagy [44]. Moreover, *p53* involves suppression of *BIRC5* at both mRNA and protein expression levels to promote apoptosis [43]. Lower doses of Vernolac extract and paclitaxel were used to maintain the cells in viable conditions until the cells were under the early apoptotic stage at the time of RNA extraction. Vernolac extract led to a significant (P < 0.001) upregulated tumor suppressor *TP53* expression, with a 9.67-fold change at 10 *μg/mL* and 13.29 fold change at 20 μg/mL compared to the untreated control (Figure 06). Concurrently, Vernolac extract-treated NTERA-2 cells resulted in significant (P < 0.05) downregulation of mammalian target of rapamycin (*mTOR*) and *BIRC5* mRNA expression at both 10 and 20 μg/mL concentrations(Figure 06). According to Abeysinghe et al. [15], it was shown that Vernolactone in *V. zeylanica* significantly induced apoptosis and autophagy in NTERA-2 cl.D1 cells through modulation of the PI3K/Akt/mTOR pathways. Key phytochemicals of *Nigella sativa*; Beta catenin, Alpha-hederin and Thymoquinone, have been reported to inhibit DNA synthesis, induce cell cycle arrest, and promote apoptosis via both p53-dependent and p53-independent mechanisms [13,33,34,45]. Collectively, the above-mentioned studies indicate that active compounds in the constituents of Vernolac extract exert their anticancer effects through coordinated modulation of apoptotic and autophagy-related pathways.

Vernolac extract comprises *V. zeylanica* aerial parts, *N. sativa* seeds, *H. indica* roots, *S. glabra* rhizome, and *L. zeylanica* aerial parts investigated in the present study, exerting anti-proliferative and anti-cancer effects via apoptosis, autophagy, and cell cycle arrest through upregulation of mRNA expression of *TP53*and downregulation of *BIRC5* and *mTOR* mRNA expression. The observed cytotoxic and apoptotic effects likely result from the action of a single active compound and/or synergistic interactions among various phytochemicals within the extract, potentially offering advantages over single-compound therapies [13,22,35,45]. Furthermore, in vitro investigations are necessary to demonstrate and confirm the precise molecular mechanisms underlying the anticancer activity, and subsequent *in vivo* studies are essential to evaluate its efficacy and the biological effects of Vernolac extract within living organisms (physiologically relevant systems), particularly its safety profile.

## Conclusion

This study demonstrates that Vernolac extract showed selective antiproliferative effects on NTERA-2 cl.D1, cancer stem-like cells by inducing apoptosis and autophagy, while exhibiting minimal cytotoxicity toward noncancerous epithelial cells (MCF-10A). These findings support the potential of Vernolac extract as a promising therapeutic candidate targeting cancer stem cells. Limitations of this study include the absence of assays such as CSC markers, autophagy flux measurements, apoptosis validation, and migration controls. These limitations will be addressed in future investigations on Vernolac.

## Acknowledgments

The authors gratefully acknowledge the support provided by the Institute of Biochemistry, Molecular Biology and Biotechnology (IBMBB), University of Colombo. The authors gratefully acknowledge Fadna Life Science Pvt. Ltd for kindly providing the Vernolac extract and supporting its chemical characterization, which greatly contributed to the advancement of this research.

## CRediT author statement

**PCR**: Investigation, Formal analysis, Writing – original draft, Writing – review and editing; **DCS, MMM**: Writing – original draft, Writing – review and editing; **SDVG, NNS, DNP**: Writing – review and editing; **UR**,**KSS**: Supervision, Writing – review and editing; **SRS**: Conceptualization, Writing – review and editing, Supervision, Project administration.

